# AI-Driven Efficient *De Novo* design of GLP-1RAs with Extended Half-Life and Enhanced Efficacy

**DOI:** 10.1101/2025.03.26.645438

**Authors:** Ting Wei, Xiaochen Cui, Jiahui Lin, Zhuoqi Zheng, Taiying Cui, Liu Cheng, Xiaoqian Lin, Junjie Zhu, Xuyang Ran, Xiaokun Hong, Zhangsheng Yu, Haifeng Chen

## Abstract

Peptide drug has revolutionized modern therapeutics, offering novel treatment avenues for various diseases. However, low efficacy, time consumption, and high cost hinder peptide drug design and discovery. We present an efficientin approach that integrates deep learning-based protein design with efficient functional screening, enabling the rapid design of biotechnologically important peptides with improved the stability and efficacy. We designed 10,000 *de novo* GLP-1 receptor agonists (GLP-1RAs), of which 60 met the stability, efficacy, and diversity criteria in the vitral functional screening. *In vitro* validation revealed a 62% success rate, while *in vivo* experiment demonstrated that two designed GLP-1RAs exhibited significantly extended half-lives, approximately three times longer than Semaglutide. In diabetic and obesity mouse models, the most competitive candidate showed superior therapeutic efficacy over Semaglutide. Our AI driven peptide design pipline integrats protein design, functional screening, and experiment validation, reducing the number of iterations required to find novel peptide candidates. The entire process, from design to screening, can be completed in a single cycle within two weeks.

## Introduction

Peptides are emerging as a crucial class of therapeutics due to their specificity, efficacy, and safety in treating various diseases, ranging from oncology, metabolic disorders, infectious diseases, and neurological conditions^1^. Peptide medications have many benefits, including low side effects, high biological activity, specificity, and effectiveness^2^. The global peptide therapeutics market size was estimated at USD 49.13 billion in 2024 and is predicted to increase from USD 52.59 billion in 2025 to approximately USD 83.75 billion by 2034 ^3^. Despite the expanding application of peptides in biomedical fields, peptides drugs face challenges such as low therapeutic efficacy, poor stability, short plasma half-life, susceptibility degradation by digestive enzymes^1,4^.

Recently, deep learning-based protein design methods have emerged as powerful tools for generating *de novo* proteins with desirable properties^5-7^, such as remarkable thermal stability^8^ and high activity^9-11^. Advances in artificial intelligence (AI) are also paving new paths for peptide drug design^12^. Unimolecular GCGR/GLP1R dual agonists model have been developed to design new GCGR/GLP1R dual agonists with enhanced receptor activation *in vitro* ^13^. HelixDiff^14^, a diffusion model for generating peptide, has designed a Glucagon-like peptide-1 (GLP-1) analogue that demonstrates *in vitro* GLP-1 receptor activation comparable to native GLP-1. A computational approach was utilized in designing peptide inhibitors that specifically target β-catenin and NF-κB^15^. AMP-Designer leveraged protein language model to design antimicrobial peptides (AMPs) with antibacterial efficacy^6^. Integraing computational design methods and a high throughput screening for generating G protein-coupled receptors (GPCRs) agonists and antagonists with high affinity, potency and selectivity^16^. However, most peptide design methods have only been validated *in silico*, with limited *in vitro* validation and almost no *in vivo* evaluation, and have not been compared with the most effective approved drugs. There remains a long way to go before designed peptides can be considered candidates for preclinical therapeutic development.

Besides design strategies, functional screening also plays a crucial role in modern drug discovery, accelerating the identification of promising candidates for experimental assays from thousands to millions of designed proteins. Deep learning-based protein design method integrated with functional screening has been successfully utilized to engineer various protein enzymes, such as serine hydrolases ^9^, myoglobin^5^, tobacco etch virus protease^11^. This two-step design pipeline has significantly improved the accuracy and efficiency of function protein design while reducing time and costs associated with traditional methods. Compared to enzyme design, peptide design presents unique challenges, as peptides are susceptible to proteolytic degradation and metabolic instability, necessitating optimization of plasma half-life and biological activity.

Here, we integrates deep learning-based protein design with efficient computational screening, enabling the successful design of promising peptide candidates with high stability, specificity, and efficacy. We firstly used ProteinMPNN^17^ for protein design, which has been successfully applied in previous studies to engineer proteins with enhanced stabilities and target-binding affinities. We then narrowed down the designed peptides for experimental assay based on stability, efficacy, and diversity screening. Stability is evaluated through enzymatic degradation, helicity, isoelectric point, hydrophobicity. Efficacy is assessed by folding ability and binding affinity with molecular dynamic (MD) simulation. The diversity aspect aims to design novel peptides that circumvent existing patent protections.

We demonstrate the effectiveness of our two-step peptide design method by designing Glucagon-like peptide-1 receptor agonists (GLP-1RAs) ^18^. The designed GLP-1RAs has enhanced biological properties, such as extended half-life, lower blood glucose, and weight loss effects in the mouse models. GLP-1RAs have gained significant attention for their efficacy in managing type 2 diabetes mellitus (T2DM) ^19,20^, obesity^19,20^, and mental health disorders.^21^. GLP-1 RA drugs have been developed from daily formulations to weekly formulations. The currently approved long-acting GLP-1RA including once weekly Exenatide^22^, Dulaglutide^23^, and Semaglutide^23^. Current research focuses on developing ultralong-acting GLP-1RA with extended half-life and better efficacy^24^. Ultralong-acting GLP-1RAs have the advantage of reduced dosing frequency, ease of use, and better safety profiles ^25-28^.

We present a two-step peptide design pipeline that combines deep leaning-based protein design with computaioal screening to efficiently design *de novo* peptides with improved activity, stability, efficacy. We employed ProteinMPNN for designing 10,000 novel GLP-1RAs, of which 60 passed the functional screening based on stability, efficacy, and diversity criteria. Notably, we successfully designed a GLP-1RA with a half-life approximately three times longer than Semaglutide and demonstrated superior efficacy compared to Semaglutide in diabetic nephropathy and obesity. Engineering new GLP-1RA with extended half-life and better efficacy requires time-consuming and expensive iterative cycle of design-make-test-analyse (DMTA)^29^. Our pipline integrates AI-powered, high-through protein design with efficient functional screening, enabling the successful design of *de novo* GLP-1RA candidates in a single cycle lasting approximately two weeks.

## Results

Fig. 1 provides an overview of the deep learning-based peptide design pipeline, integrating efficient functional screening and experimental validation to enable the rapid design of promising peptide candidates with extended half-life and enhanced efficacy. Using GLP-1RAs as an example, within two weeks, we successfully designing 60 *de novo* GLP-1RAs with improved stability, specificity, and efficacy. Among them, three were validated *in vivo*, revealing a half-life approximately three times longer than Semaglutide and improved efficacy in glucose-lowering and weight loss. The entire process, from design to validation, was concluded in one cycle.

**Fig. 1.**
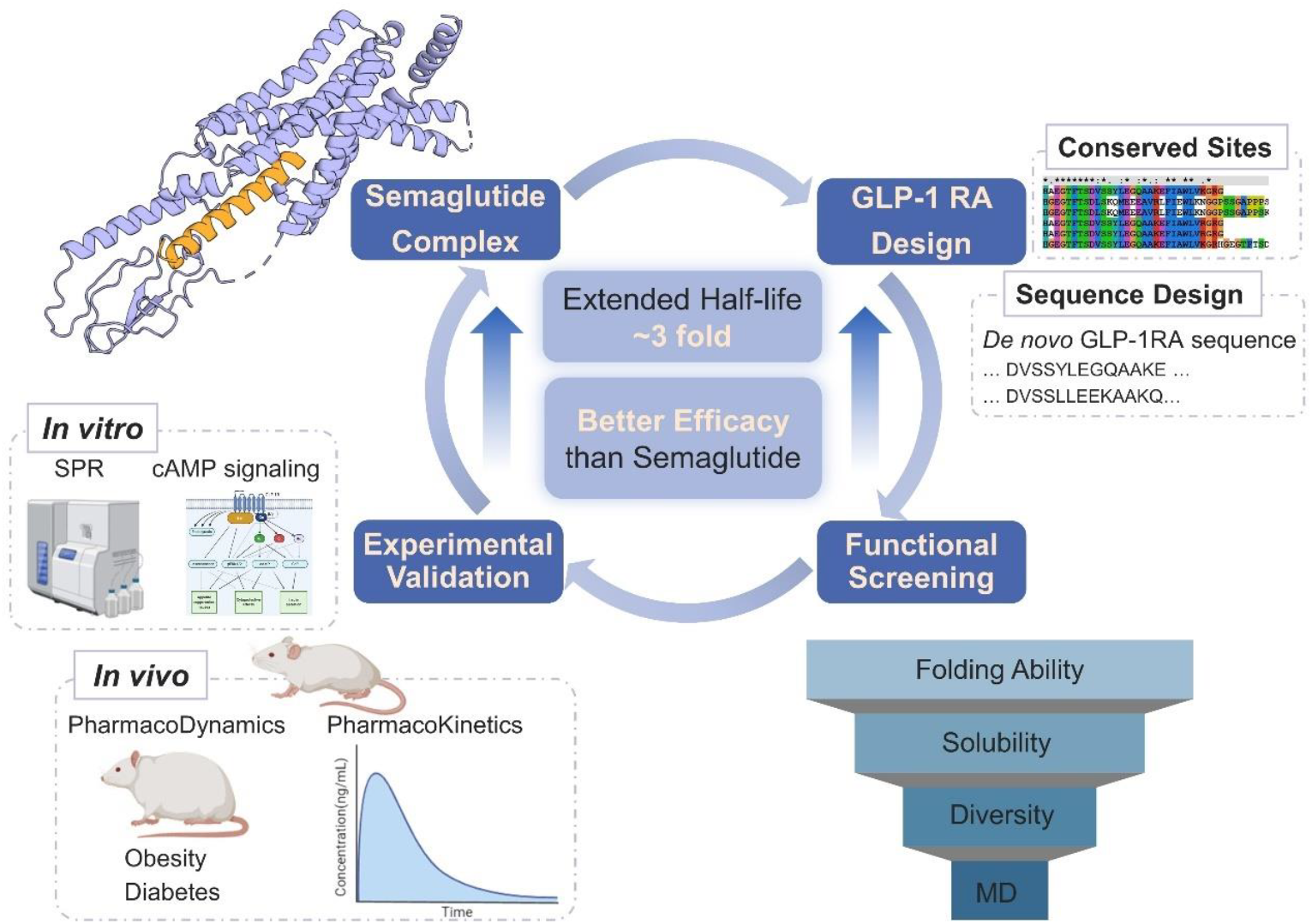
The deep learning-based peptide design-efficient functional screening-experiment validation pipeline. This workflow includes *de novo* GLP-1RAs design, computational screening, and experimental validation through *in vitro* and *in vivo* studies.

### *De novo* GLP-1RAs Design

Our *de novo* GLP-1 RAs design pipeline consists of two primary stages: conserved sites analysis and ProteinMPNN *de novo* sequences design. Before designing the GLP-1RAs, we defined conserved sites or hotspot residues of GLP-1RAs as critical functions for target recognition, binding, and activation of the GLP-1 receptor (GLP-1R). 12 residue points (7H, 8Aib, 9E, 10G, 11T, 12F, 13T, 14S, 15D, 17S, 26K, 34R, 37G) were fixed according to conservation analysis. These residues interact with GLP-1R’s transmembrane core or binding to the extracellular domain of the GLP-1R.

We fixed these conserved sites and then used ProteinMPNN to produce functional designs of GLP-1RA. The experimentally determined crystal structure of the Semaglutide-GLP-1R complex (PDB 7KI0) as a template for our design. Using ProteinMPNN, we generated 10,000 *de novo* GLP-1RAs sequences by fixing 12 highly conserved sites. ProteinMPNN, a deep leaning method for protein complex design, has been successfully deployed in previous studies to engineer protein, such as ubiquitin^10^, protein nanomaterials^7^, myoglobin^5^, TEV protease^19^. In this study, we extend its application to peptide design.

### Functional Screening

High-throughput computational screening is an efficient method for narrowing down designed peptides, indentifying promising drug candiates with desirable properties and accelerating drug discovery process. The 10, 000 designed GLP-1Ras underwent virtual screening based on stability, efficacy, and diversity sciteria. Finally, 60 with disiralbe properties, including extended biological half-life, high binding affinity, and greater diversity compared to approved GLP-1RAs, were selected for *in vitro* and *in vivo* validation (Fig. 2A and Fig. S1-2).

**Fig. 2.**
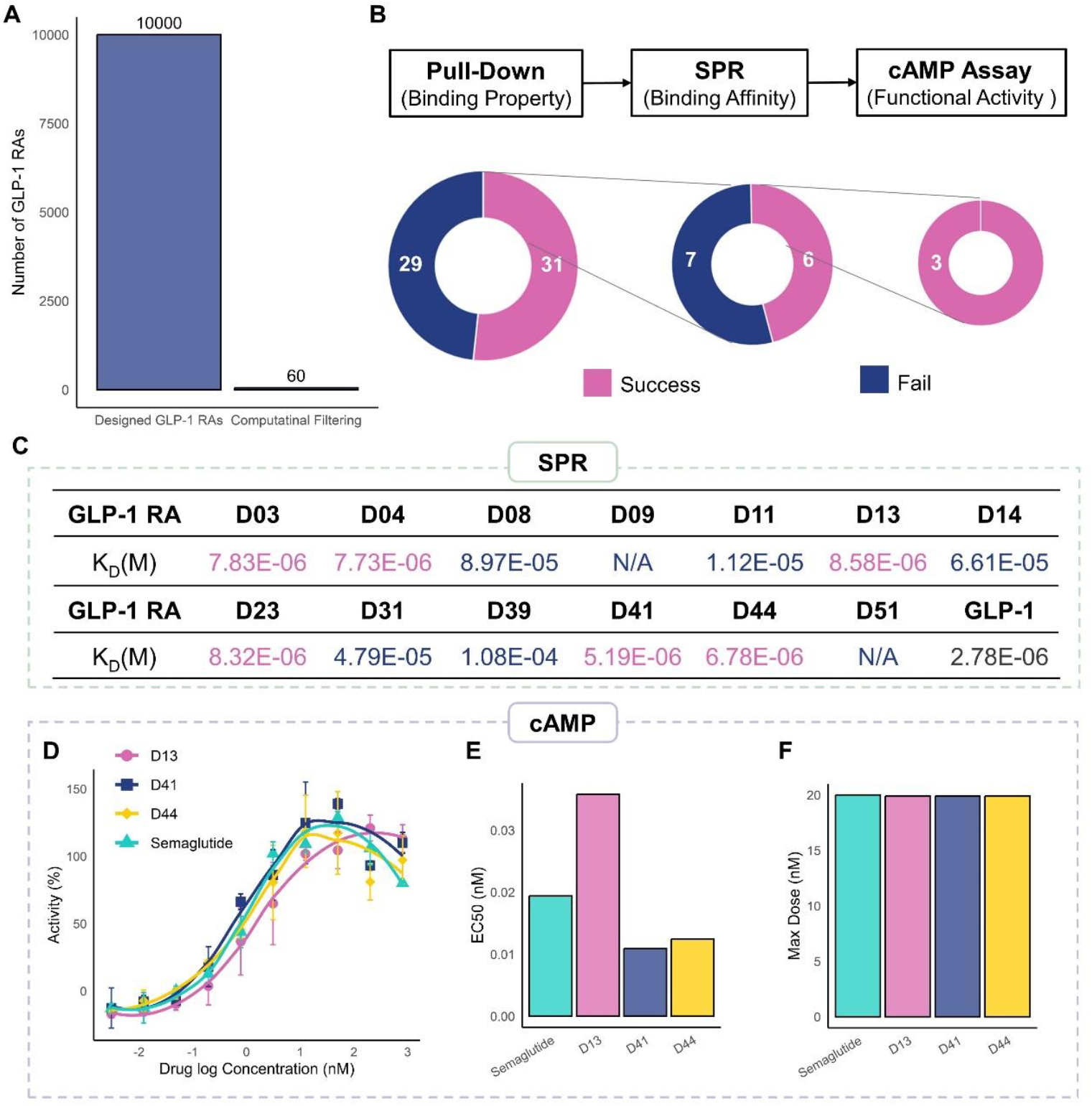
*In vitro* Functional Screening of GLP-1RAs. **(A)** The number of GLP-1RAs after computational screenings. (B) The number of successful and failed sequences at each *in vitro* screening step: GST pulldown assay, SPR, and cAMP accumulation assay. (C) Binding affinity of GLP-1RAs assessed by SPR. (D) the cAMP dose-response curves for GLP-1RAs were measured in HEK-293 cells expressing the cloned human GLP-1R. Data represent means ± SD of two independent experiments performed in ten concentrations. (E) the 50% activity concentration (EC_50_). (F) the Max concentration (Emax).

**Fig. 3.**
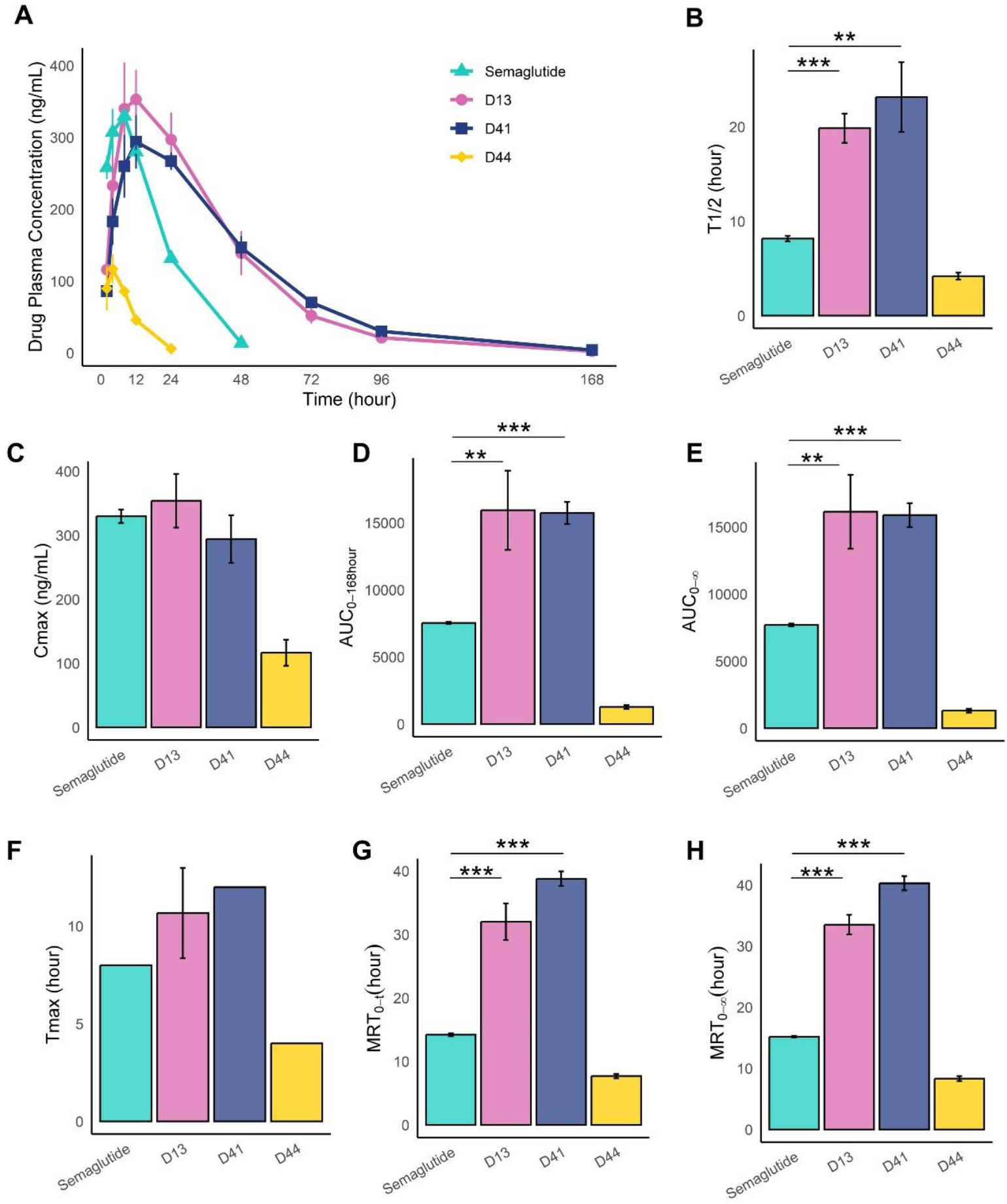
The PK result in rates (Mean±SD, n=3). (A) The drug plasma concentration-time curve for Semaglutide (0.05 mg/kg), D13 (0.05 mg/kg), D41 (0.05 mg/kg), and D44 (0.05 mg/kg). (B) half-life (T_1/2_). (C) The maximum plasma concentration (C_max_). (D-E) The area under the concentration-time curve of AUC_0-168 hours_ and AUC_0-∞_. (F) Time to Maximum Concentration (T_max_). (G-H) The mean residence time of MRT_0-168hours_ and MRT_0-∞_. *P < 0.05, **P < 0.01, ***P < 0.001.

Peptides drugs are susceptibility degradation by digestive enzymes, resulting in poor stability and a short plasma half-life. Studies indicated that NEP-24.11 can cleave GLP-1RAs at six potential cleavage sites in the central and C-terminal regions^30^. To improve stability and extend the half-life of the designed GLP-1RAs, sequences containing these cleavage sites were filtered out. Additionally, factors such as helicity, isoelectric point, hydrophobicity impact the peptide potency^13,31^, we further filter the designed GLP-1RAs based-on the net charge, hydrophobicity and spatial aggregation propensity (SAP) score (details in the Methods).

Binding ability of peptides to their receptor is closedly associated with efficacy, and was evaluated using the folding ability of complex structure and MD simulations. AlphaFold2 was used to predict the complex structures of the designed GLP-1RA sequences and GLP-1R. The folding capability of the designed GLP-1RAs were assessed using several key parameters: The predicted Local Distance Difference Test (pLDDT) scores, which evaluate the quality of the predicted protein structure; the TM-score and Root Mean Square Deviation (RMSD), which assess how well the designed structure aligns with the native structure; and the interface pAE, which evaluates the confidence in the interactions at the interface between GLP-1RA and GLP-1R in the complex. MD simulations provided insights into the binding affinity, which measures the binding strength between GLP-1RA and GLP-1R.

The diversity focuses on designing novel peptides that circumvent existing patent protections. Phylogenetic trees were generated to categorize sequences, and a single representative sequence was selected in each phylogenetic cluster. Finally, a set of 60 unique GLP-1RA sequences was selected for subsequent experimental evaluation.

### *In vitro* Experimental Validation

We performed an *in vitro* functional screening of 60 unique GLP-1RA sequences using a combination of GST pulldown assay, surface plasmon resonance (SPR), and intracellular cAMP (cyclic adenosine monophosphate) accumulation assay (Fig. 2B and Fig. S2). The GST pulldown assay served as an effective preliminary screening, revealing 31 sequences that could bind to GLP-1R, Subsequent SPR analysis further evaluated the binding affinity of these sequences, with 6 GLP-1RAs demonstrating binding affinities comparable to Semaglutide. Finally, the cAMP assay assessed the functional activation of GLP-1R, revealing that three GLP-1RAs (D13, D41 and D44) could activate cAMP signaling. The number of successful and failed GLP-1RAs at each screening step is summarized in Fig. 2B.

#### The GST pulldown assay

this assay was employed to evaluate the binding capability of GST-peptide fusion proteins to GLP-1R, providing a qualitative measure of receptor interaction. GST and GST-GLP-1 were employed as negative control and positive control respectively. Serving as a preliminary screening step, the assay identified 31 GLP-1RAs capable of binding to GLP-1R, while the remaining 29 sequences failed to show interaction (Fig. 2B).

#### SPR

SPR was utilized as the next screening step to quantitatively assess the binding affinity between the GST-fusion peptides and GLP-1R. The dissociation constant (K_D_) for GLP-1 (Semaglutide peptide) binding to GLP-1R was 2.78 × 10^−6^ M. Of the 31 GLP-1RAs that passed the GST pulldown assay, 13 underwent SPR analysis (Fig. 2B). Among these, 6 GLP-1RAs (D03, D04, D13, D23, D41, and D44) exhibited binding affinities comparable to GLP-1, with K_D_ values on the order of E-06 (Fig. 2C).

#### cAMP accumulation assay

We further evaluated the functional activity of the GLP-1RAs by measuring their ability to elicit cAMP signaling, a key downstream response of GLP-1R activation. Among the six GLP-1RAs that passed the SPR analysis, three (D13, D41 and D44) underwent cAMP accumulation assay (Fig. 2D). D41 and D44 exhibited superior cAMP signaling, with half-maximal effective concentrations (EC_50_) of 0.011 nM and 0.012 nM, respectively, compared to Semaglutide’s 0.019 nM (Fig. 2E). D13, with an EC_50_ of 0.036 nM, demonstrated similar cAMP signaling. The max dose (Emax) was in the narrow range of 19.89-20.00 nM (Fig. 2F). These results indicated that all three GLP-1RAs could effectively activate intracellular cAMP signaling.

GST pulldown assay, SPR assay, and intracellular cAMP accumulation assay identified three GLP-1RA candidates (D13, D41, D44) with strong receptor binding and effective functional activity, making them suitable for further *in vivo* experimental evaluation.

### Extended half-life of GLP-1RAs in Pharmacokinetics (PK)

Pharmacokinetics (PK) refers to the study of a drug’s absorption, distribution, metabolism, and excretion (ADME) within the body. Among the three GLP-1RA candidates, D41 and D44 exhibited significantly longer half-lives compared to Semaglutide in SD rat. Specifically, D13 exhibited a half-life (T_1/2_) of 19.86 hours, approximately 2.43 times longer than Semaglutide’s 8.17 hours. Similarly, D41 displayed a prolonged T_1/2_ of 23.16 hours, 2.83 times that of Semaglutide (Fig. 4A and 4B, and Table 1).

**Fig. 4.**
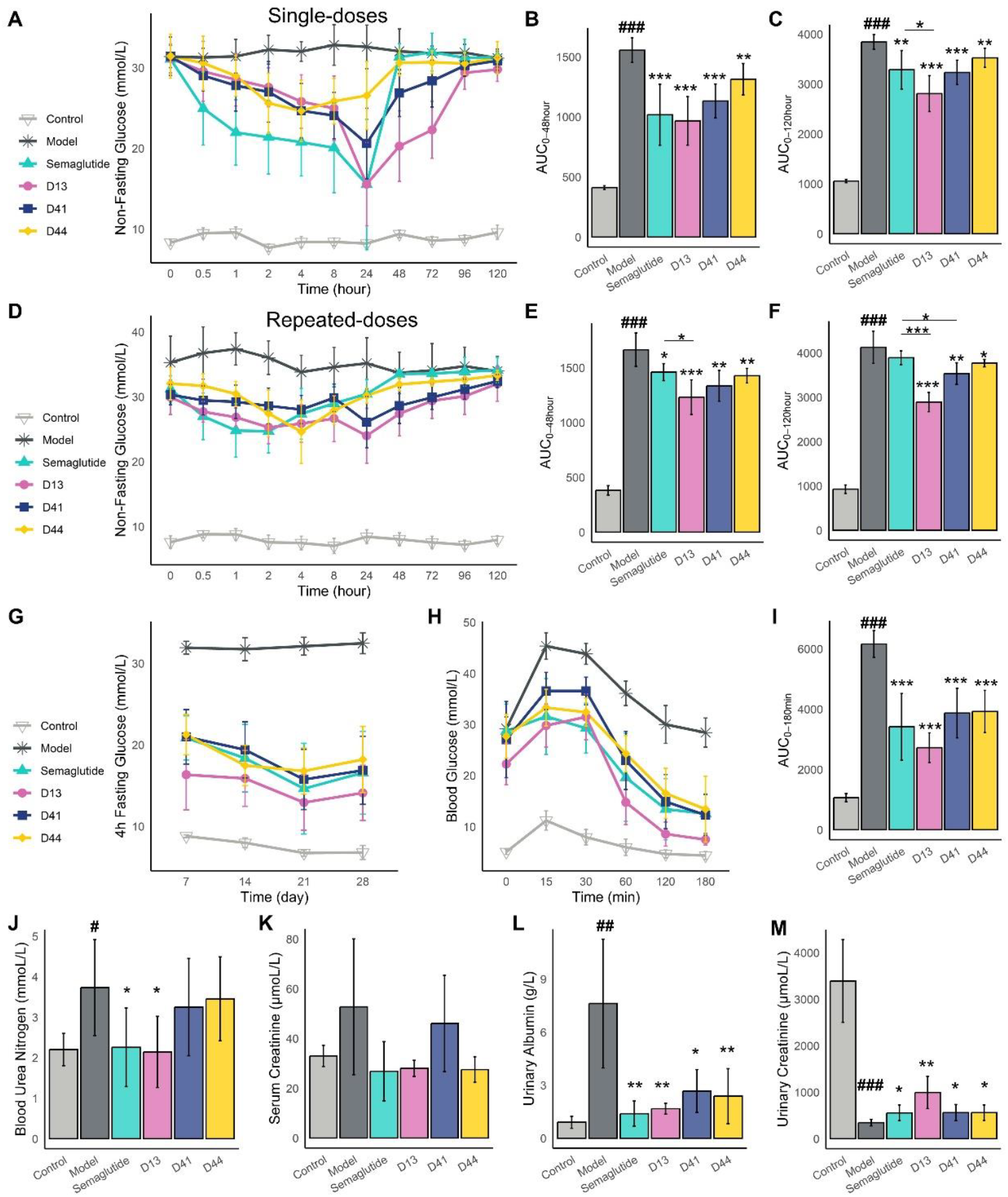
Glucose-lowering of GLP-1RAs in Diabetic model (Mean±SD, n=6). The db/db mice treated with GLP-1RAs and Semaglutide at 10 nmol/kg for single dose (A-C) and multiple doses for 28 days (D-M). (A) the non-fasting glucose-time curve after single dose. (B) the area under the non-fasting glucose-time curve (AUC) after single dose within 0-48 hour (AUC_0-48 hour_) and (C) within 0-120 hour (AUC_0-120 hour_). (D) the non-fasting glucose-time curve after multiple doses. (E) the area under the non-fasting glucose-time curve after multiple doses within 0-48 hour (AUC_0-48 hour_) and (F) within 0-120 hour (AUC_0-120 hour_). (G) 4 hour fasting blood glucose during 28-days continuous administration period. (H) the intraperitoneal glucose tolerance test (iPGTT) after multiple doses. (I) the area under the iPGTT curve (AUC) with 0-180min after multiple doses. (J-M) kidney injury biomarkers of blood urea nitrogen (BUN), serum creatinine (Scr), and urinary albumin levels, and urinary creatinine levels after multiple administration. # Compared with normal group, *compared with model group. #P < 0.05, ##P < 0.01, ###P < 0.001, *P < 0.05, **P < 0.01, ***P < 0.001.

D13 also showed a maximum plasma concentration (C_max_) of 353.96 ng/mL, which is higher than Semaglutide’s C_max_ of 329.82 ng/mL (Fig. 4C). The area under the concentration-time curve (AUC_0-t_), representing the total drug exposure over 0-168 hours, was 15939.88 and 15737.19 h*ng/mL for D13 and D41, respectively, both approximately 2.1 times than Semaglutide’s 7538.16 h*ng/mL (Fig. 4D). AUC_0-∞_, representing the total drug exposure over 0-infinite time, was also higher for D13 and D41 compared to Semaglutide (Fig. 4E).

Time to Maximum Concentration (T_max_) was 10.67 hour for D13 and 23.16 hour for D41, both longer than Semaglutide’s 8.17 hour (Fig. 4F). The mean residence time (MRT_0-t_), indicating the average duration for drug remains in the body before elimination within 0-168 hours, was 32.02 for D13 and 38.79 hour for D41, approximately 2.3 times and 2.7 times that of Semaglutide (14.21 hour) (Fig. 4G). Similarly, MRT_0-∞_, representing the average duration for drug remains in the body before complete elimination, was higher for both D13 and D41 compared to Semaglutide (Fig. 4H).

These results highlight the extended half-life and higher plasma concentration of D13 than Semaglutide, indicating enhanced stability and the potential for prolonged therapeutic effects compared to Semaglutide.

### Lower Blood Glucose Levels of GLP-1RAs in Diabetic Nephropathy

The hypoglycemic effect of the GLP-1RAs was evaluated in diabetic db/db mouse model. GLP-1RAs were administrated subcutaneously both as a single-dose and as multiple doses for 28 days (10 nmol/kg administered once daily). The hypoglycemic effect of single dose focused on acute reduction in blood glucose levels. This analysis provides valuable insights into the time-effect relationship, which is compared with the PK results. The hypoglycemic effect of multiple doses primarily examine the long-term effects of glucose-lowering and cumulative impacts.

### Acute glucose-lowering of single-dose GLP-1RAs

We evaluate the acute glucose-lowering of single dose D13, D41, and D44 in the diabetic db/db mouse model. Non-fasting glucose were monitored at timepoints of 0, 0.5, 1, 2, 4, 8, 24, 48, 72, 96, 120 hours. We observed that D13, D41, D44, and Semaglutide elicited comparable glucose-lowering effects compared with model group (Fig. 5A-C).

**Fig. 5.**
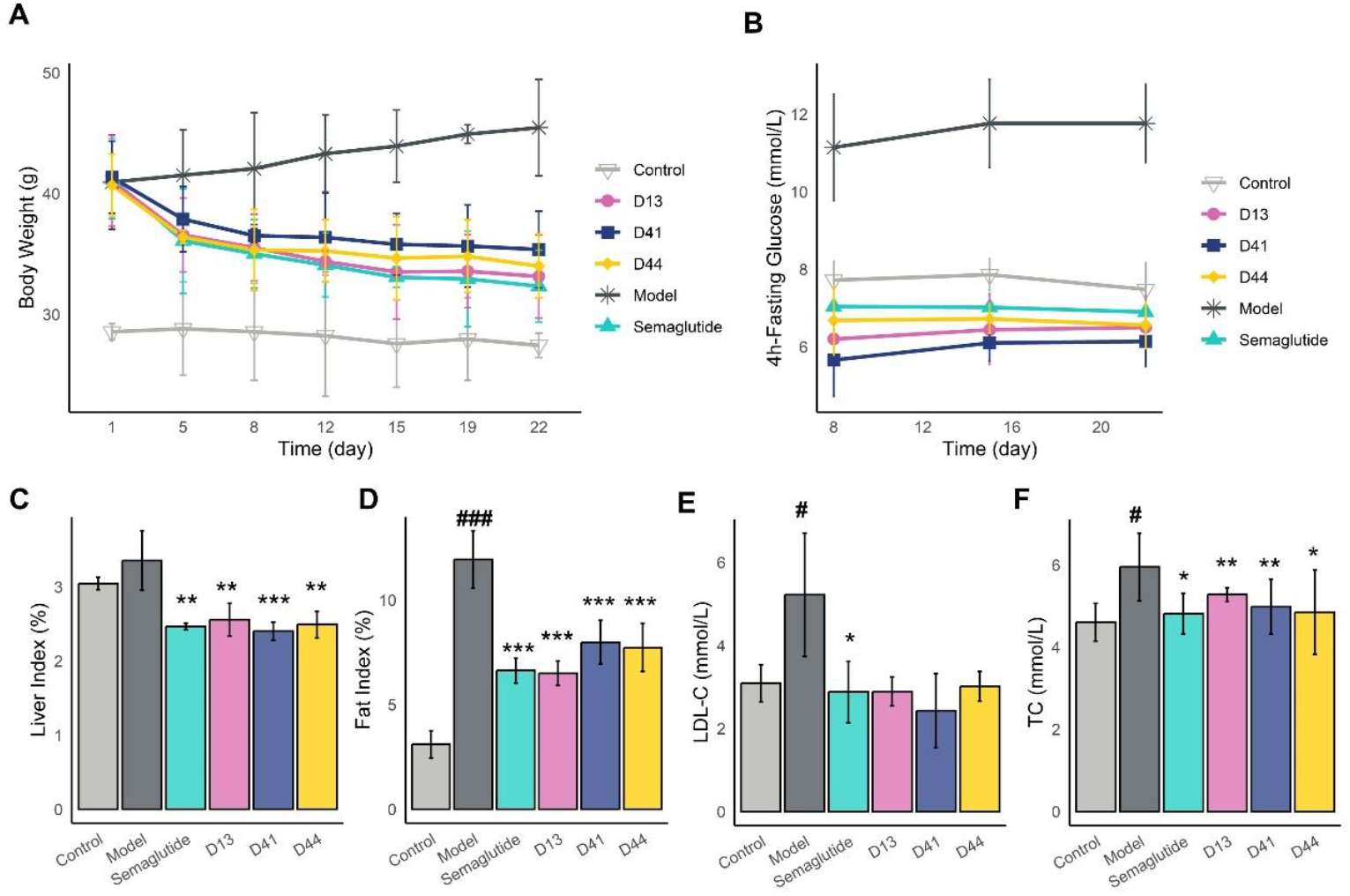
Weight loss of GLP-1RAs in Obesity model (Mean±SD, n=5). The DIO mice treated with GLP-1RAs and Semaglutide from 0 to 22 days at 10 nmol/kg, measuring (A) body weight and (B) 4 hour fasting blood glucose. (C) liver index. (D) fat index. (E) serum LDL-C. (F) serum TC. # Compared with normal group, *compared with model group, #P < 0.05, ##P < 0.01, ###P < 0.001, *P < 0.05, **P < 0.01, ***P < 0.001.

In the Semaglutide (10 nmol/kg) group, non-fasting blood glucose levels began to decrease at 0.5 hours after administration and remained reduced until 24 hours. Blood glucose levels reached their lowest point between 8 and 24 hours and returned to baseline (model group levels) by 48 hours (Fig. 5A and Fig. 5B). In the D13 group, non-fasting blood glucose levels began decreasing at 0.5 hours post-administration and persisted for up to 96 hours. Blood glucose levels reached their lowest point between 24 and 72 hours, returning to near baseline levels by 96 hours (Fig. 5A and Fig. 5C). The lowest blood glucose level in the D13 group (15.53±8.06 mmol/L) was comparable to that of the Semaglutide group (15.57±5.16 mmol/L). Notably, non-fasting glucose levels after 24 hours administration in the D13 group was significantly reduced compared with the Semaglutide group at three times points, with P-values of 0.0002, 0.0002, 0.1694, 0.0436 at 48, 72, 96, 120 hours, respectively. The D41 group has the similar effect with D13 group, blood glucose levels reached their lowest point between 8 and 48 hours (20.6±4.37 mmol/L), returning to near baseline levels by 96 hours.

These findings were consistent with PK, suggest that D13 may offer prolonged glucose control compared to Semaglutide, making them potential candidates for sustained management of blood glucose in type 2 diabetes.

### Long-term effects of glucose-lowering after multiple doses GLP-1RAs

To evaluate the long-term glucose-lowering, D13, D41, and D44 were administered subcutaneously over 28 days period (every day for 28 days) in the diabetic db/db mouse model. At end of the 28 days, non-fasting glucose were monitored at timepoints of 0, 0.5, 1, 2, 4, 8, 24, 48, 72, 96, 120 hours. In the D13 group (10 nmol/kg), non-fasting blood glucose levels in db/db mice began decreasing at 0.5 hours post-administration and persisted for up to 96 hours. The lowest blood glucose level in the D13 group (24.00±4.24mmol/L) was lower than that of the Semaglutide group (24.68±3.34 mmol/L). Notably, the blood glucose after 8 hours in the D13 group was significantly reduced compared with the Semaglutide group at four times points, with P-values of 0.0081, 0.0016, 0.0047, 0.028 at 24, 48, 72, 96 hours, respectively. (Fig. 4D-F). This may be attributed to the dose accumulation effect of D13, given its longer half-life and higher C_max_. A similar effect was observed in the D41 group, where the blood glucose was significantly lower than the Semaglutide group after 8 hours, with P-values of 0.04, 0.0017, 0.0012, 0.04 at 24, 48, 72, 96 hours, respectively (Fig. 4D-F).

Over a 28-days continuous administration period, 4 hour-fasting blood glucose levels were measured once a week. The 4 hour-fasting blood glucose of D13 group were also lower than those in the Semaglutide group, with no statistical significance (Fig. 4G).

The intraperitoneal glucose tolerance test (iPGTT) were performed 120 hours after multiple doses. Blood glucose was measured at 10, 30, 60, 120, and 180 min after glucose loading. Semaglutide, D13, D41, and D44 all demonstrated improved glucose tolerance during 0-180 minutes (Fig. 4H). D13 exhibited slightly greater efficacy than Semaglutide, though the difference was not statistically significant. The AUC_0-180min_ for D13 (2725.75±491.03 mmol/L*min) was lower than that of Semaglutide’s 3414±1105.75 mmol/L*min, indicating enhanced glucose tolerance (Fig. 4I).

Further investigation into renal protection revealed that Semaglutide and D13 significantly reduced markers of kidney injury in *db/db* mice, including blood urea nitrogen (BUN), serum creatinine (Scr), and urinary albumin levels, while concurrently increasing urinary creatinine levels (Fig. 4J-M).

### Weight Loss of GLP-1RAs in Obesity

To assess the weight loss efficacy of GLP-1RAs, we treated diet-induced obese (DIO) mice with multiple administrations over 21 days. DIO mice exhibited a decrease in body weight during treatment with D13, D41, and D44. D13, D41, and D44 resulted in body weight reductions of 19.25%, 14.42%, and 16.46%, respectively, while semaglutide led to a 21.54% reduction (Fig. 5A). The weight loss efficacy of D13 was comparable to that of semaglutide, with no significant difference between the two treatments. These data demonstrate the effectiveness of D13 in reducing body weight in DIO mice.

The 4 hour fasting blood glucose in DIO mice decreased in Semaglutide, D13, D41, and D44 group, with D13 and D44 showing slightly lower levels than the Semaglutide group, though the difference was not statistically significant (P-value >0.5) (Fig. 5B). Semaglutide, D13, D41, and D44 group exhibited reduced liver and fat index (Fig. 5C-D). Serum LDL-C in DIO mice were significantly reduced in the Semaglutide, D13, D41, and D44 (Fig. 5E). While serum TC showed a decreasing trend in the Semaglutide, D13, D41, and D44, the difference was not statistically significant (Fig. 5F).

### Sequence and Structure analysis of GLP-1RAs

To provide sequence and structural insights into the GLP-1RAs with longer half-life and improved efficacy, we performed a comprehensive analysis of sequence, structure, and evolutionary relationships. Hoping to offer additional design principles for developing more effective GLP-1RAs with prolonged half-life and improved efficacy.

We first examined sequence differences between successful and failed GLP-1RAs in GST pulldown assays and SPR assay. Interestingly, no significant differences between successful and failed groups were observed in the recovery and diversity distributions. These metrics, which are critical in AI-based protein sequence design, suggest that factors beyond sequence recovery and diversity influence the success of GLP-1RAs. (Fig. 6A).

**Fig. 6.**
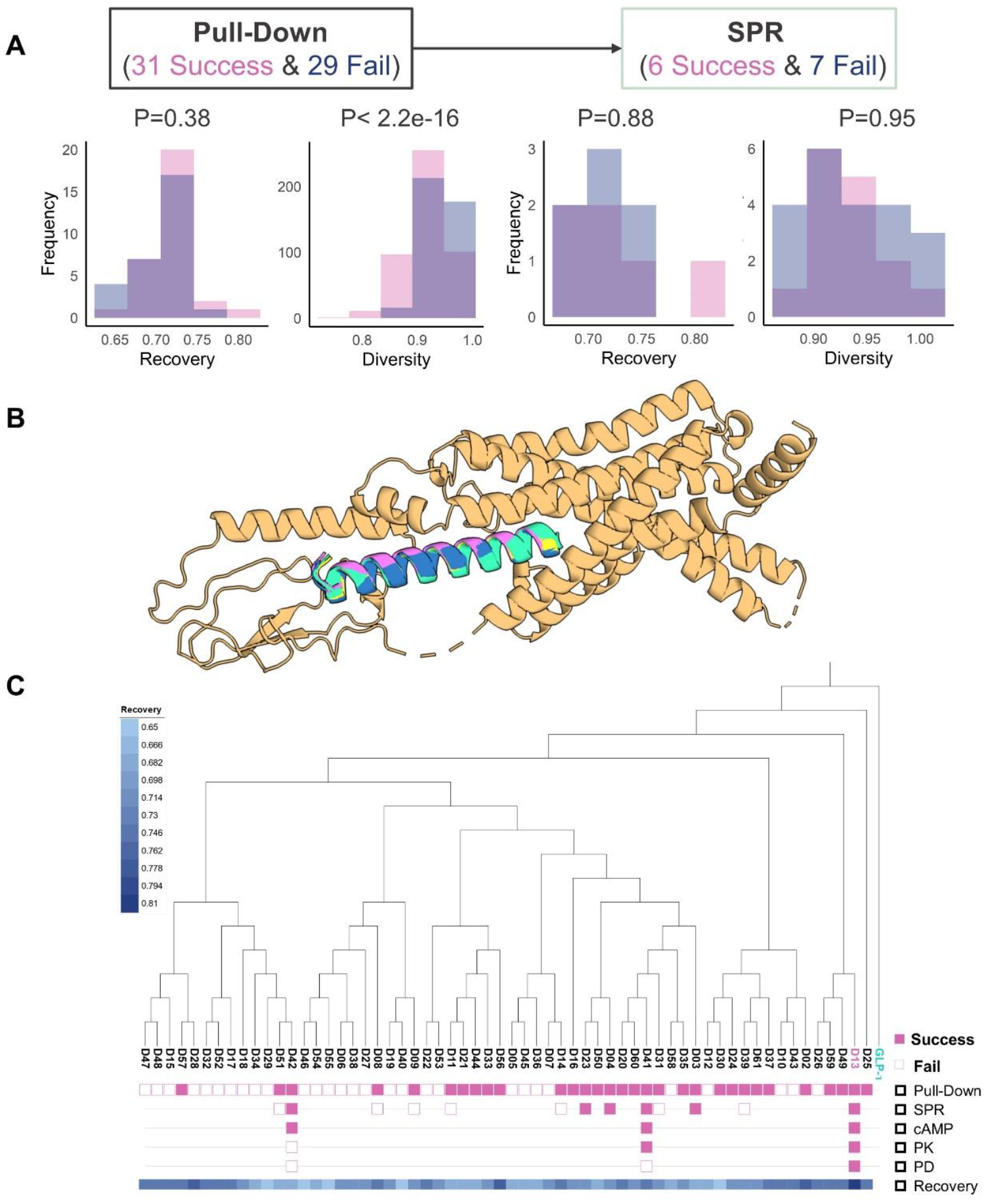
Sequence, structural, and evolutionary analyses of GLP-1RAs. (A) Sequence analysis: Comparison of recovery and diversity distributions between successful and failed GLP-1RAs using GST pulldown and SPR assays. (B) Structural analysis: AlphaFold2-predicted complex structure of GLP-1RA and GLP-1R. (C) Evolutionary analysis: Phylogenetic tree of 60 GLP-1RAs constructed to explore evolutionary relationships.

To further explore the structural basis of GLP-1RA, we used AlphaFold2 to predict the complex structure of GLP-1RA and GLP-1R (Fig. 6B). The predicated structures revealed that the GLP-1RAs have a conserved α-helix structure, necessary for GLP-1R activate. These findings highlight the importance of maintaining this structural feature in the design of novel GLP-1RAs.

Furthermore, to understand the evolutionary relationships among GLP-1RAs, we constructed a phylogenetic tree comprising 60 GLP-1RAs (Fig. 6C and Fig. S4). Both successful and failed GLP-1RAs were broadly distributed throughout the tree, indicating no distinct evolutionary clustering between the two groups. Notably, the root of the phylogenetic tree is Semaglutide. D13, which exhibits a longer half-life and superior efficacy, is positioned close to Semaglutide in the tree, suggesting a shared evolutionary lineage and sequence similarity (recovery = 0.81). This close proximity supports the hypothesis that D13 is an ancestral GLP-1RA, similar in origin and mechanism to Semaglutide. Additionally, D25 (recovery = 0.74), another GLP-1RA not yet experimentally validated, is also positioned near Semaglutide. Although D25 was not included in this round of experimental validation, its sequence recovery and structural features make it a promising candidate for further investigation.

## Discussion

Peptides have emerged as a unique drug class for treating a wild range of disease, including diabetes, cancer, and central nervous system disorders. However, designing biotechnologically essential peptides required expensive, time-consuming, and extensive validation. Deep learning-based protein design methods has proven to be a powerful tool for generating novel proteins with desired properties^32^. Here, we introduce a deep learning-based peptide design pipelines that combine deep learning protein sequence design and efficient funcational screening, enabling the successful design of novel GLP-1RAs with extended half-life and enhanced properties efficacy compared to Semaglutide. For GLP-1RAs design, 10,000 designed sequences were subsequently subjected to funcational screening based on stability, efficacy, and diversity, resulting in a much-reduced number of peptide candidates, about 60 GLP-1RAs for experimental evaluation.

The GST pulldown assay identified 31 GLP-1RAs that interact with GLP-1R. From these, we selected 13 candidates for the SPR assay, and 6 exhibited binding affinities comparable to Semaglutide. We then chose 3 GLP-1RAs (D13, D41, and D44) for cAMP assay. The EC_50_ rank was: D41 (0.011 nM) > D44 (0.012 nM) > Semaglutide (0.019 nM) > D13 (0.036 nM). As only a subset of candidates was validated, additional GLP-1RAs may also be capable of activating the cAMP second messenger signaling pathway.

In Pharmacokinetics (PK), both D13 and D41 exhibited significantly improved pharmacokinetic properties relative to Semaglutide, including a longer half-life, extended T_max_, and enhanced C_max_. These improvements suggest that D13 and D41 may offer prolonged therapeutic effects. The half-life (T_1/2_) ranking was: D41 (23.16 hours) > D13 (19.86 hours) > Semaglutide (8.17 hours) > D44 (4.19 hours). The C_max_ ranking was: D13 (353.96 ng/mL) > Semaglutide (329.82 ng/mL) > D44 (116.60 ng/mL) > D44 (116.60 ng/mL). Notably, the *in vivo* findings were not consistent with the cAMP assay results. In PK, the D44 was worst.

In diabetic mice with simple administration and multiple administration, the hypoglycemic effects of D13 and D41 were notably prolonged compared to Semaglutide. These findings are consistent with the observed pharmacokinetic data, further supporting that extended half-life contributes to more durable blood glucose control. D13 demonstrated significantly lower blood glucose levels than Semaglutide in both single-dose and multiple-dose regimens. Weight reduction with D13 was slightly lower than Semaglutide, but the difference was not statistically significant. Efficacy outcomes varied across disease models, indicating that factors such as disease subtype might influence treatment response of the same drug. This highlights the importance of designing GLP-1RAs candidates tailored for specific diseases.

Our AI-powered high-through protein design-functional screening-experiment validation pipelines provides a promising way for improving the stability and efficacy of biotechnologically import peptide drug. This piplien has been evaluated only in GLP-1RAs. We are now designing GLP-1R/GCGR/GIPR triple agonists and hope that the *in vitro* experiment will surpass the performance of Retatrutide^33^.

## Materials and Methods

### Design *De novo* GLP-1RAs

For conserved site analysis, we collected the sequences of GLP-1RAs marked drug, including GLP-1, Exenatide, Lixisenatide, Liraglutide, Semaglutide, and Albiglutide. Clustal X was used to identify highly conserved sites of GLP-1RAs in multiple sequence alignments (Fig. S3).

The conserved sites and crystal structure of the Semaglutide-GLP-1R complex (PDB 7KI0) was used as inputs for ProteinMPNN. To generate *de novo* GLP-1RAs, with the GLP-1R chain being fixed and the Semaglutide chain being selected for optimized sequences. We used ProteinMPNN to generated 10,000 seuqnces using the default parameters, including sampling temperature=0.1.

### Computational Screening of designed GLP-1RAs

#### Stability

Using ProteinMPNN, we generated a total of 10,000 GLP-1RA sequences. Semaglutide, the initial template for our design, consists 31 amino acids. Of these, 13 conserved residues were fixed, while the remaining 18 amino acids were designed by ProteinMPNN. Due to the limited design space and conserved structural constraints (α-helix), a substantial number of repetitive sequences were generated during this process. After eliminating redundant sequences, 1,443 unique sequences were obtained.

*In vitro* studies revealed that the enzyme NEP-24.11 can cleave GLP-1 at six potential sites in the central and C-terminal regions, with particular vulnerability between Glu27-Phe28 and Trp31-Leu32, along with additional cleavage sites at Asp15-Val16, Ser18-Tyr19, Tyr19-Leu20, and Phe28-Ile29. To improve stability and extend the half-life of the designed GLP-1RAs, we filtered out sequences containing these cleavage sites.

The net charge, hydrophobicity and SAP were used to estimate the protein solubility. The net charge of a protein plays a crucial role in its solubility. SAP was calculated for each residue based on a combination of solvent accessibility and hydrophobicity. Hydrophobicity primarily controls the exposure of non-polar residues on the protein surface.

#### Efficacy

AlphaFold2 was employed to predict the complex structures of designed GLP-1RA sequences and GLP-1R. The best rank structure of each complex was further calculated TM-score, RMSD, pLDDT, and interface pAE. TM-score between the designed complex structure (GLP-1 RA and GLP-1R) and the native complex structure (Semaglutide and GLP-1R); RMSD between the designed GLP-1RA structure and Semaglutide native structures; pLDDT for the designed GLP-1RA structures; The average pAE of interchain residue pairs (interface pAE) of GLP-1 RA and GLP-1R complex. The critical thresholds for these parameters were derived from the native complex structure between Semaglutide and GLP-1R.

ff03CMAP force field was used for MD simulation, and this force field was developed by our group. The solvent model used was TIP4P-Ew, a model proved to be suitable for ordered protein. The antechamber was used to parameterize the ligand molecule. Firstly, energy minimization, heating and equilibrium of the system were carried out. The energy of the system was minimized by the steepest descent method of 3000 steps and the conjugate gradient method of 3000 steps. After energy minimization, the system was heated from 0 to 321 K in a time of 50 ps and then performs an energy balance of 100 ps at constant pressure and temperature of 321 K. In the whole process, the long-range electrostatic interaction was calculated by PME algorithm, and the covalent bonds of all hydrogen atoms were constrained by SHAKE algorithm. The cut-off value for the van der Waals interaction and the short-range electrostatic interaction was set at 8 Å. The final simulation process was carried out at NPT and temperature of 321 K, and the simulation time was 20 ns.

#### Diversity

Phylogenetic trees were generated using the Neighbor-Joining method in MEGA and visualized with iTOL (v6). To ensure sequence diversity, a single representative sequence was selected from each phylogenetic cluster.

### Study Design of *in vitro* Experiment

#### GST pulldown assay

DNA coding sequences corresponding to 60 GLP-1RAs and Semaglutdie were cloned into PGEX-4T-1 (Cytiva, GE Healthcare) between BamHI and XhoI, respectively. Constructs were transformed into *E. coli* BL21 (DE3) for GST-peptide fusion protein expression. Human GLP1R protein (MedChemExpress LLC, HY-P700468) were added to each GST-fusion protein coupled beads. Proteins binding to beads were eluted by 1% sodium dodecyl sulphonate. Samples were loaded on SDS-PAGE and results were displayed by coomassi blue staining. GST protein and GST-wild type peptide were used as negative control and positive control respectively.

#### Surface plasmon resonance (SPR)

Binding affinity between GLP1R and GST-fusion peptides were investigated by surface plasmon resonance on Biacore 8k system (Cytiva). Briefly, the CM5 sensor chip (Cytiva, 29149603) was used to measure the binding kinetics between GLP1R and GST-fused peptides. Affinity K_D_ values were fitted by steady state affinity fitting models (1:1 binding model) using Biacore Insight Evaluation Software, experimental data were analyzed using the Biacore Insight Evaluation Software, with concentration as the X-axis and steady-state response as the Y-axis. Purified GST protein was also loaded for kinetic analyses as a negative control.

#### Human GLP-1R cell cAMP assay

Intracellular cAMP accumulation was measured using the cAMP detection kit (Cisbio, 62AM4PEJ) based on the homogeneous time-resolved fluorescence (HTRF) technology. Briefly, HEK293 cells stably expressing GLP-1R were transferred to OptiPlate-384 plate. Add GLP-1RAs and Semaglutide were diluted with PBS buffer supplemented with 500 µM, and were incubated with cells for 30 min at room temperature. Time-resolved FRET signals were measured on an EnVision (PerkinElmer) at 665 nm and 620 nm. All the dose-response curve fits were analyzed with R using an equation of log (GLP-1RAs) vs activity.

### Study Design of in vivo Experiment

#### Study Design of Pharmacokinetics (PK)

SD male rats were administered a single dose of 0.05 mg/kg of Semaglutide (Ozempic^®^) and the GLP-1RAs. Blood samples were collected at 10 post-administration time points: 2h, 4h, 8h, 12h, 24h, 48h, 72h, 96h, and 168h. Plasma samples were analyzed using LC-MS/MS, and pharmacokinetic parameters such as T_1/2_, C_max_, T_max_, AUC_0-t_, and MRT_0-t_ were calculated using WinNonlin.

### Study Design of db/db Diabetic Nephropathy models

#### Acute glucose-lowering after single administration

Male db/db mice were randomly divided into five groups, with six mice per group: model group, Semaglutide group (10 nmol/kg), D13 group (10 nmol/kg), D41 group (10 nmol/kg), and D44 group (10 nmol/kg). Additionally, six db/m mice were included as the control group. Each animal was administered a single dose of the drug. Non-fasting blood glucose levels were monitored at multiple time points post-administration: 0 h, 0.5 h, 1 h, 2 h, 4 h, 8 h, 24 h, 48 h, 72 h, 96 h, and 120 h.

#### long-term effects of glucose-lowering after single administration

Two weeks after completing the single-dose administration study, once the fasting blood glucose of the db/db mice (measured after 4 hours of fasting) returned to the level of the model group, a 4-week continuous administration was conducted. During the continuous administration period, fasting blood glucose levels were measured once a week. On day 28, non-fasting blood glucose levels were measured at 0 h, 0.5 h, 1 h, 2 h, 4 h, 8 h, 24 h, 48 h. Afterward, the intraperitoneal glucose tolerance test (iPGTT) was performed after overnight fasting. Urine samples were collected for the analysis of urine albumin and urine creatinine content. The next day (after overnight fasting), animals were deeply anesthetized with CO2, and blood samples were collected for the measurement of glycated hemoglobin, creatinine, and blood urea nitrogen levels. Both kidneys were excised, weighed, and the kidney to body weight ratio was calculated.

#### Study Design of DIO Obesity Models

Male C57BL/6J mice were fed a high-fat diet (D12492) for approximately 10 weeks. The control group and model group were administered PBS, while the Semaglutide group (10 nmol/kg) and D13 group (10 nmol/kg), D41 group (10 nmol/kg), and D44 group (10 nmol/kg) received continuous administration for 4-week. During the continuous administration period, body weight was monitored twice per week. After the administration, the mice were deeply anesthetized with CO2, and blood was collected to measure blood lipids, including serum total cholesterol (TC) and low-density lipoprotein cholesterol (LDL-C). The abdominal cavity was then opened, and white adipose tissue from the epididymis, abdominal wall, and mesentery was collected, weighed, and the coefficient of fat was calculated. The liver was excised, weighed, and it’s the coefficient of liver was also calculated.

### Statistical Analysis

Statistical analyses were conducted using R (v4.3.1). Data, error bars, sample size/replicates, and statistical tests used are defined in the Fig. legends. In general, data were analyzed with the two-sided t-test.

## Reference

1 Muttenthaler, M., King, G. F., Adams, D. J. & Alewood, P. F. Trends in peptide drug discovery. Nature reviews Drug discovery 20, 309–325 (2021).

2 Chen, Z., Wang, R., Guo, J. & Wang, X. The role and future prospects of artificial intelligence algorithms in peptide drug development. Biomedicine & Pharmacotherapy 175, 116709 (2024).

3 Pereira, A. J., de Campos, L. J., Xing, H. & Conda-Sheridan, M. Peptide-based therapeutics: challenges and solutions. Medicinal Chemistry Research 33, 1275–1280 (2024).

4 Otvos, L. vol. 19 869–872 (Taylor & Francis, 2024).

5 Sumida, K. H. et al. Improving protein expression, stability, and function with ProteinMPNN. Journal of the American Chemical Society 146, 2054–2061 (2024).

6 Wang, J. et al. Discovery of antimicrobial peptides with notable antibacterial potency by an LLM-based foundation model. Science Advances 11, eads8932 (2025).

7 de Haas, R. J. et al. Rapid and automated design of two-component protein nanomaterials using ProteinMPNN. Proceedings of the National Academy of Sciences 121, e2314646121 (2024).

8 Vázquez Torres, S. et al. De novo designed proteins neutralize lethal snake venom toxins. Nature, 1–7 (2025).

9 Lauko, A. et al. Computational design of serine hydrolases. Science, eadu2454 (2025).

10 Kao, H.-W. et al. Robust Design of Effective Allosteric Activators for Rsp5 E3 ligase using the machine learning tool ProteinMPNN. ACS Synthetic Biology 12, 2310–2319 (2023).

11 Yeh, A. H.-W. et al. De novo design of luciferases using deep learning. Nature 614, 774–780 (2023).

12 Goles, M. et al. Peptide-based drug discovery through artificial intelligence: towards an autonomous design of therapeutic peptides. Briefings in Bioinformatics 25, bbae275 (2024).

13 Puszkarska, A. M. et al. Machine learning designs new GCGR/GLP-1R dual agonists with enhanced biological potency. Nature Chemistry 16, 1436–1444 (2024).

14 Xie, X., Valiente, P. A., Kim, J. & Kim, P. M. HelixDiff, a Score-Based Diffusion Model for Generating All-Atom α-Helical Structures. ACS Central Science 10, 1001–1011 (2024).

15 Chen, S. et al. Design of target specific peptide inhibitors using generative deep learning and molecular dynamics simulations. Nature Communications 15, 1611 (2024).

16 Muratspahić, E. et al. De novo design of miniprotein agonists and antagonists targeting G protein-coupled receptors. bioRxiv, 2025.2003.2023.644666, doi:10.1101/2025.03.23.644666 (2025).

17 Dauparas, J. et al. Robust deep learning–based protein sequence design using ProteinMPNN. A Science 378, 49–56 (2022).

18 Wang, L. et al. Therapeutic peptides: current applications and future directions. Signal transduction and targeted therapy 7, 48 (2022).

19 Zheng, Z. et al. Glucagon-like peptide-1 receptor: mechanisms and advances in therapy. Signal Transduction and Targeted Therapy 9, 234 (2024).

20 Trujillo, J. M., Nuffer, W. & Smith, B. A. GLP-1 receptor agonists: an updated review of head-to-head clinical studies. Therapeutic advances in endocrinology and metabolism 12, 2042018821997320 (2021).

21 De Giorgi, R. et al. An analysis on the role of glucagon-like peptide-1 receptor agonists in cognitive and mental health disorders. Nature Mental Health, 1-20 (2025).

22 Knop, F. K., Brønden, A. & Vilsbøll, T. Exenatide: pharmacokinetics, clinical use, and future directions. Expert opinion on pharmacotherapy 18, 555–571 (2017).

23 Jendle, J. et al. Efficacy and safety of dulaglutide in the treatment of type 2 diabetes: a comprehensive review of the dulaglutide clinical data focusing on the AWARD phase 3 clinical trial program. Diabetes/metabolism research and reviews 32, 776–790 (2016).

24 Schneider, E. L., Hangasky, J. A., Ashley, G. W. & Santi, D. V. Long-Acting, Longer-Acting, and Ultralong-Acting Antiobesity Peptides. Journal of Medicinal Chemistry (2025).

25 Zhu, D. et al. Efficacy and safety of GLP-1 analog ecnoglutide in adults with type 2 diabetes: a randomized, double-blind, placebo-controlled phase 2 trial. Nature Communications 15, 8408 (2024).

26 Liu, Y. et al. The safety, tolerability, pharmacokinetics and pharmacodynamics of GZR18 in healthy American and Chinese adult subjects. Diabetes, Obesity and Metabolism.

27 Marechal, A. et al. 1674-P: Development of a Once-a-Month Formulation of Semaglutide from an Innovative Injectable and Biodegradable Hydrogel. Diabetes 73 (2024).

28 Zhang, Y. et al. 119-OR: Safety, Tolerability, Pharmacokinetics, and Pharmacodynamics of a Novel Ultra-Long-Acting GLP-1 Receptor Agonist (ZT002) in Healthy Subjects. Diabetes 73, 119–OR (2024).

29 Ghiandoni, G. M., Evertsson, E., Riley, D. J., Tyrchan, C. & Rathi, P. C. Augmenting DMTA using predictive AI modelling at AstraZeneca. Drug discovery today 29, 103945 (2024).

30 Plamboeck, A., Holst, J., Carr, R. & Deacon, C. Neutral endopeptidase 24.11 and dipeptidyl peptidase IV are both mediators of the degradation of glucagon-like peptide 1 in the anaesthetised pig. Diabetologia 48, 1882–1890 (2005).

31 Chabenne, J. et al. A glucagon analog chemically stabilized for immediate treatment of life-threatening hypoglycemia. Molecular metabolism 3, 293–300 (2014).

32 Zhang, K. et al. Artificial intelligence in drug development. Nature Medicine 31, 45–59, doi:10.1038/s41591-024-03434-4 (2025).

33 Jastreboff, A. M. et al. Triple–hormone-receptor agonist retatrutide for obesity—a phase 2 trial. New England Journal of Medicine 389, 514–526 (2023).

